# KRAS inhibition is an effective therapy for appendiceal adenocarcinoma

**DOI:** 10.64898/2026.04.07.717107

**Authors:** Saikat Chowdhury, Ichiaki Ito, Vinay K Pattalachinti, Abdelrahman MG Yousef, Mahmoud MG Yousef, Sacha El Khoury, Nicholas Hornstein, Ashlee Nichole Seldomridge, David Hong, Micheal J. Overman, Melissa W. Taggart, Wai Chin Foo, Beth Helmink, Keith F. Fournier, John Paul Shen

**Affiliations:** Department of Gastrointestinal Medical Oncology, The University of Texas MD Anderson Cancer Center, Houston, TX, USA; Northwell Cancer Institute, New York, NY, USA; Department of Surgical Oncology, University of Texas MD Anderson Cancer Center, Houston, TX, USA; Department of Investigational Cancer Therapeutics, The University of Texas MD Anderson Cancer Center, Houston, TX, USA; Department of Pathology, Division of Pathology–Lab Medicine, University of Texas MD Anderson Cancer Center, Houston, TX, USA

## Abstract

**Background:** Appendiceal adenocarcinoma (AA) is a rare cancer with limited treatment options. *KRAS* is the most commonly mutated gene in AA and a promising therapeutic target, but its preclinical and translational relevance in AA remains unclear.

**Methods:** We evaluated KRAS^G12D^–specific (MRTX1133) and pan-KRAS inhibitor (RMC-6236) in *KRAS^mut^* organoid and orthotopic PDX models of AA. Tumor-intrinsic and microenvironmental responses were characterized using multi-omics profiling. Clinical outcomes were also assessed in six heavily pre-treated AA patients treated with KRAS inhibitors.

**Results:** MRTX1133 was highly effective for KRAS^G12D^ organoids (IC50=4.1 nM); both KRAS^G12D^ and KRAS^G12V^ organoids were sensitive to RMC-6236 (IC50=4.4 nM vs 0.5 nM, respectively). In orthotopic PDX models of peritoneal carcinomatosis from AA, MRTX1133 significantly reduced tumor growth in the KRAS^G12D^ model TM00351, and RMC-6236 reduced tumor growth in KRAS^G12V^ model AAPDX-16. Pathologic evaluation showed dramatically reduced tumor cellularity, proliferation, and pERK expression as well as induction of apoptosis. Gene Sets Enrichment Analysis (GSEA) revealed significant downregulations of ‘E2F targets (NES=-1.9, p-adj=0.06) and the newly developed ‘RAS/ERK (NES=-2.3, p-adj=0.06)’ gene set, consistent with the observed decrease in cell proliferation. There was marked upregulation of EMT (NES=2.7, FDR<0.001) and TGF-β signaling (NES=2.3, FDR=0.004) in remaining tumor cells, suggesting these pathways could confer resistance. scRNA-seq analysis of TME showed dramatic shifts in cancer-associated fibroblasts (CAFs), with KRAS inhibition driving a shift from normal fibroblasts to inflammatory CAFs, and upregulation of interferon alpha and gamma pathways, suggesting that KRAS inhibition can activate innate immune response in the setting of peritoneal metastases. In a cohort of 6 heavily pre-treated patients with AA treated with KRAS inhibitors (1 G12D, 3 G12C, 2 pan-KRAS), all had biochemical response based on CEA/Ca19-9 or ctDNA and clinical benefit by RECIST criteria (1 CR, 1 PR, 4 SD).

**Conclusions:** While effective suppression of RAS/ERK signaling by KRAS inhibitors reduces tumor growth, adaptive activation of EMT and TGF-β pathways may mediate resistance in *KRAS^mut^* AA. Additionally, KRAS inhibition remodels TME and may enhance innate immune signaling. These findings support continued clinical development of KRAS inhibitors in AA and provide a rationale for combination strategies targeting resistance pathways and stromal remodeling.

## Background

Appendiceal adenocarcinoma (AA) is a rare, orphan disease with no systemic therapy currently approved by the United States FDA specifically for use in appendiceal cancer. Historically, chemotherapy developed for colorectal cancer (CRC) was used to treat patients with AA; however, recent studies have demonstrated that AA is distinct from CRC at a molecular level (*1*) and that CRC chemotherapy is ineffective for many patients with AA(*2*). *KRAS* is the most frequently mutated gene in AA, occurring in nearly 80% of mucinous adenocarcinomas and roughly 65% of colonic-type adenocarcinomas (*1, 3*). Long considered ‘undruggable’, multiple small-molecule inhibitors of KRAS have been developed, including both pan-KRAS and allele-specific (G12C & G12D) inhibitors (*4*). This breakthrough paved the way for successful clinical trials of two KRAS^G12C^ inhibitors, AMG510 (sotorasib) and MRTX849 (adagrasib), for the treatment of KRAS^G12C^-mutant solid tumors (*5*). Subsequently, these drugs received FDA approval to treatment refractory metastatic CRC and were incorporated into the NCCN guidelines (*6, 7*). Two additional single-agent drugs, divarasib, and olomorasib, have been developed to target KRAS^G12C^-mutant solid tumors, demonstrating efficacy in clinical trials for non-small cell lung cancer (NSCLC) and mCRC (*8, 9*). Several other KRAS^G12C^-targeting drugs, including JNJ-74699157, IBI351, HBI-2438, JDQ443, JAB-21822, and BPI-421286, are currently in the preclinical phase or entering early stages of clinical trials (5). A recently developed non-covalent inhibitor, MRTX1133, which specifically targets KRAS^G12D^, has demonstrated remarkable efficacy in preclinical studies and has entered phase I/II clinical trials (10, 11). Other KRAS^G12D^ inhibitors in development include RMC-9805, HRS-4642, TH-Z835, BI-2852, JAB-22000, and ERAS-4 (12). Moreover, ongoing efforts aim to develop single-agent drugs that inhibit multiple KRAS alleles, such as pan-KRAS inhibitors like RMC-6236 and BI-2865, which hold immense potential to revolutionize targeted therapy for solid tumors with *KRAS^Mut^* (13, 14).

Single-agent KRAS-G12C inhibitors, Sotorasib and Adagrasib, have demonstrated impressive objective response rates (ORR) in patients with Non-small cell lung cancer (NSCLC) and moderate response in patients with metastatic CRC, with phase II trials reporting ORRs of NSCLC: 37.1% vs. CRC: 9.7% for Sotorasib, and NSCLC: 42.9% vs. CRC: 23% for Adagrasib, respectively. In pancreatic ductal adenocarcinoma (PDAC), a prototypical KRAS-driven malignancy in which >90% of tumors harbor activating KRAS mutations (*15*), early clinical trials have demonstrated that direct KRAS inhibition can yield meaningful antitumor activity. In the phase I/II CodeBreaK 100 study, the KRAS(G12C) inhibitor sotorasib achieved an objective response rate (ORR) of 21% in previously treated PDAC, with a median progression-free survival (PFS) of 4.0 months (*16*). Similarly, adagrasib demonstrated objective responses in 33.3% of PDAC patients enrolled in the KRYSTAL-1 trial, further validating that pharmacologic KRAS suppression can translate into tumor regression in PDAC (*17*). More recently, a translational study reported modest responses in patients with advanced solid tumors, including pancreatic adenocarcinoma, treated with RMC-6236, providing early clinical evidence supporting this multi-allelic KRAS inhibition strategy(*18*).

These advances in PDAC, CRC, and NSCLC are highly relevant to appendiceal adenocarcinoma (AA), which also exhibits similarly high KRAS mutation frequencies yet remains a therapeutically underserved disease. Together, the clinical success of KRAS-directed therapies in these cancers and the high prevalence of KRAS mutations in AA provide a strong rationale for evaluating KRAS inhibitors as a targeted therapeutic strategy in appendiceal adenocarcinoma, a disease with a critical unmet need for effective systemic treatments. Here we report the results of the first 6 AA patients treated with KRAS inhibitors as well as results of a G12D-specific (MRTX1133) and a pan-KRAS (RMC-6236) inhibitor tested in novel *KRAS* mutant organoid and PDX models and profiled with high-content imaging, IHC, RNAseq, and Reverse Phase Proteomic Arrays (RPPA).

## Results

### KRAS is the primary oncogenic driver in Appendiceal Adenocarcinoma

Using comprehensive somatic mutation data from cBioPortal, we assessed *KRAS* alteration frequencies across a broad spectrum of tumor histologies to contextualize its role in appendiceal cancer. In mucinous appendiceal adenocarcinomas, *KRAS* mutations are strikingly pervasive, occurring in approximately 79.3% of cases (n = 130), and remain highly prevalent in colonic-type appendiceal adenocarcinomas (67.6%, n = 25) (**Figure 1A**). This high mutational frequency firmly establishes *KRAS* as the key oncogenic driver in appendiceal malignancies. When compared across major gastrointestinal (GI) cancers, the dependency on *KRAS* in appendiceal tumors parallels that of pancreatic neoplasms—pancreatic ductal adenocarcinoma exhibits *KRAS* alterations in ∼89.7% of cases (n = 2,425). By contrast, conventional colorectal adenocarcinoma harbors *KRAS* mutations in 42% of tumors (n = 1471). These comparative data underscore a spectrum of *KRAS* reliance across GI tumors, with appendiceal and pancreatic cancers clustering at the high-frequency end of the continuum (**Figure 1A**). Intriguingly, goblet cell adenocarcinoma of the appendix deviates markedly from this pattern, with a *KRAS* mutation rate of just 8.3% (n = 6). This low incidence of *KRAS* mutation suggests that goblet-cell–predominant appendiceal tumors follow an alternative oncogenic trajectory with implications for targeted therapy selection. The frequency as well as distribution of *KRAS* mutant alleles in metastatic AA was similar in our institution’s cohort (The University of Texas MD Anderson Cancer Center; MDACC) and the previously published Memorial Sloan Kettering Cancer Center (MSKCC) cohort(*3*) were highly concordant with G12D as the predominant allele in both populations (47.4% in MDACC, n = 465; 50.9% in MSKCC, n = 254, **Figure 1B**). G12V was the second most frequent allele (27.8%, 23.0%), followed by G13D (7.7%, 11.2%) and G12C (3.3%, 6.2%) in the MDACC vs MSKCC cohorts, respectively. These similarities persisted across histological subtypes—mucinous, colonic-type, and goblet cell—with G12C mutations observed exclusively in mucinous tumors in both cohorts (**supplemental figure 1A and 1B**).

**Figure 1:**
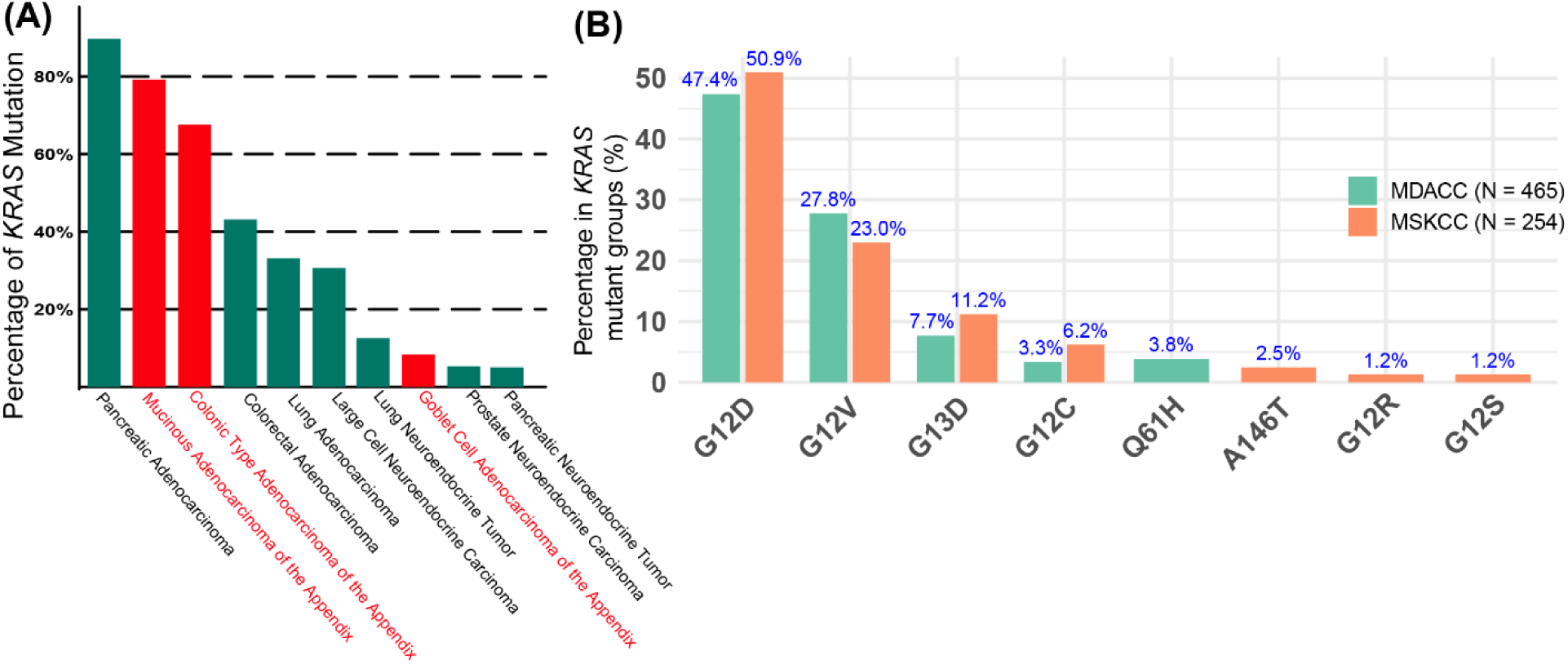
KRAS is the dominant oncogene in appendiceal cancer. (A) *KRAS* gene mutation rates across different cancer histology types. (B) Mutation frequencies of different *KRAS* alleles in *KRAS^Mut^*groups of MDACC (N = 465) and MSKCC (N = 254) cohorts.

### KRAS inhibition is highly effective in organoid models of AA

Given the predominance of activating *KRAS* variants in AA, we assessed the therapeutic potential of KRAS inhibition using organoid models. Two organoid lines were derived from PDX models; AAPDO-01 derived from a high-grade mucinous adenocarcinoma harboring *KRAS^G12D^*, *TP53^P72R^*, and *TP53^V274F^* mutations, and AAPDO-16, derived from a high-grade signet ring adenocarcinoma harboring *KRAS^G12V^*and *TP53^R282W^* mutations (**Figure 2A**). AAPDO-01 was quite sensitive to treatment with the G12D selective inhibitor MRTX1133, IC_50_ = 4.05 nM for AAPDO-01; as expected, AAPDO-16 with *KRAS^G12V^* mutation was resistant (IC_50_ =1.79 μM, **Figure 2B**). Conversely, both models were susceptible to the pan-KRAS inhibitor RMC-6236, with AAPDO-16 showing sub-nanomolar sensitivity (IC50 = 0.5 nM) and AAPDO-01 showing nanomolar sensitivity (IC50 = 4.43 nM; **Figure 2C**). KRAS inhibition induced apoptosis in sensitive organoids, consistent with a cytotoxic rather than cytostatic mechanism (**Figure 2D**). Compared with KRASG12D mutant CRC or PDAC cell models, AAPDO-01 was more sensitive to MRTX1133, as indicated by comparison of previously reported IC50 values (**Figure 2E**) (*19*). Similarly, both AA organoid models were more sensitive to RMC-6236 than both G12V and G12D models from various cancer types (**Figure 2F**) (*18*). In aggregate, these findings suggest that AA organoids are sensitive to KRAS inhibition.

**Figure 2:**
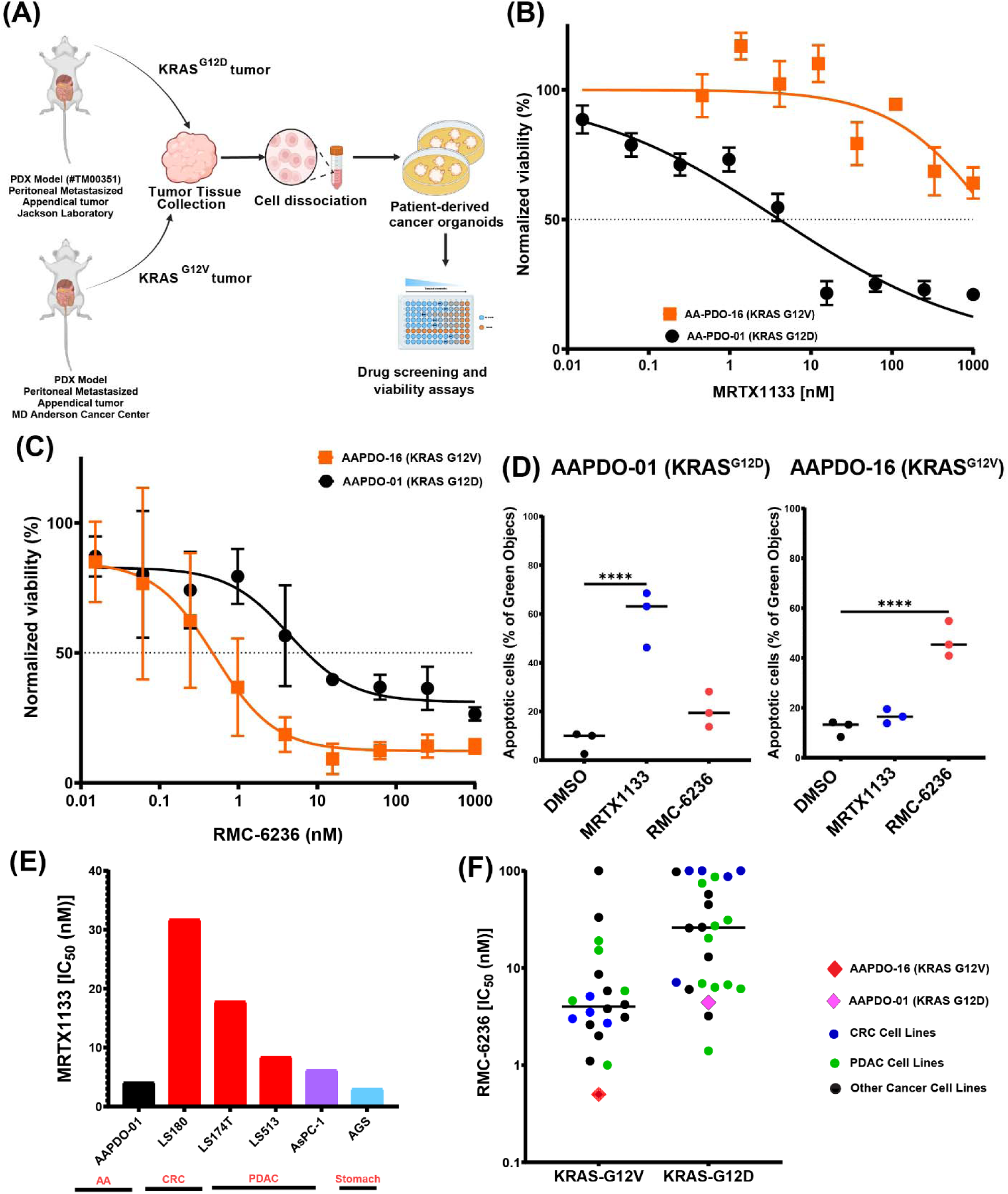
KRAS inhibition decreases the viability of AA organoids. (A) Overview of the drug screening workflow conducted in two organoid models of appendiceal adenocarcinoma (AA) tumors harboring *KRAS^G12D^* and *KRAS^G12V^* mutations. (B) MRTX1133 demonstrates greater potency in *KRAS^G12D^* organoids compared to *KRAS^G12V^*organoids. (C) RMC-6236 was effective against both *KRAS^G12D^* and *KRAS^G12V^* mutant AA organoids. (D) Apoptosis assays for MRTX1133 and RMC-6236 in both organoid models. Green object counts indicate the percentage of apoptotic or dead cells. Comparisons of IC50 values we observed for (E) MRTX1133 and (F) RMC-6236 in AA models vs previously reported IC50 values of other cancer cell lines (*18, 19*).

### KRAS inhibition is highly effective in KRAS mutant orthotopic AA PDX models

Next, we tested KRAS inhibitors in two orthotopic PDX models of AA, AAPDX-01 and AAPDX-16. For MRTX1133 treatment in the AAPDX-01 model, fresh AA tumors harboring the *KRAS^G12D^*mutation were implanted into the peritoneal cavity of NSG mice. The mice were then treated with IP-delivered MRTX1133 (15 mg/kg, BID) starting 1 month after implantation for 4 weeks, and then sacrificed for pathological evaluation of tumors (**Figure 3A**). The AAPDX-01 tumor was clearly visualized by MRI within the peritoneal cavity. MRTX1133 treatment significantly reduced tumor size relative to solvent control after 2 weeks of treatment, with the magnitude of difference increasing at 4 weeks (31.2% reduction versus vehicle control at 4 weeks, p = 0.0001; **Figure 3B, C**). Importantly, MRTX1133 did not significantly reduce body weight in mice after treatment (**Supplementary Figure 2A**). H&E staining of control tumors after 4 weeks showed densely packed, highly cellular tumors consistent with aggressive tumor growth. In contrast, tumors treated with MRTX1133 had substantially reduced tumor cellularity and showed increased stromal and immune cell infiltration (**Figure 3D**). The replacement of tumor with stroma was confirmed with Ku80 (a marker of human cells), CDX2 (a marker of intestinal, in this case, tumor cells), and vimentin staining (**Figure 3D**). Similarly, the fraction of Ki-67-positive tumor cells was reduced by MRTX1133 treatment, and the fraction of Caspase-3 tumor cells was increased (**Figure 3D**, **Supplementary Figure 2B and 2C**). Tumor cells also showed decreased or absent phosphorylated ERK (pERK) staining, consistent with effective suppression of MAPK signaling. Interestingly, pERK was increased in stromal cells, suggesting a rebound activation of ERK signaling in fibroblasts. In a second experiment, AAPDX-01 mice were treated until death or morbidity requiring euthanasia. MRTX1133 treatment significantly extended the survival of AAPDX-01 mice relative to control (median survival 35 days versus 145 days for MRTX1133-treated mice, log-rank test *P* = 0.0011, and hazard ratio 13.4, **Figure 3E**). For RMC-6236 treatment in the AAPDX-16 model, fresh AA tumors with KRAS^G12V^ mutation were implanted into the peritoneal cavity of NSG mice, which developed detectable tumors after 2 weeks (**Figure 3F**). RMC-6236 treatment (20 mg/kg IP daily, starting 2 weeks after implantation) significantly reduced tumor volume after only 7 days (52.5% reduction versus control, *p* = 0.01, **Figure 3G**) but did not significantly reduce the body weight of mice (**Supplementary Figure 2D**). Furthermore, similar to the G12D inhibitor, IHC analysis after RMC-6236 treatment showed a decreased proportion of tumor cells and a marked decrease in p-ERK (**Supplementary Figure 2E**).

**Figure 3:**
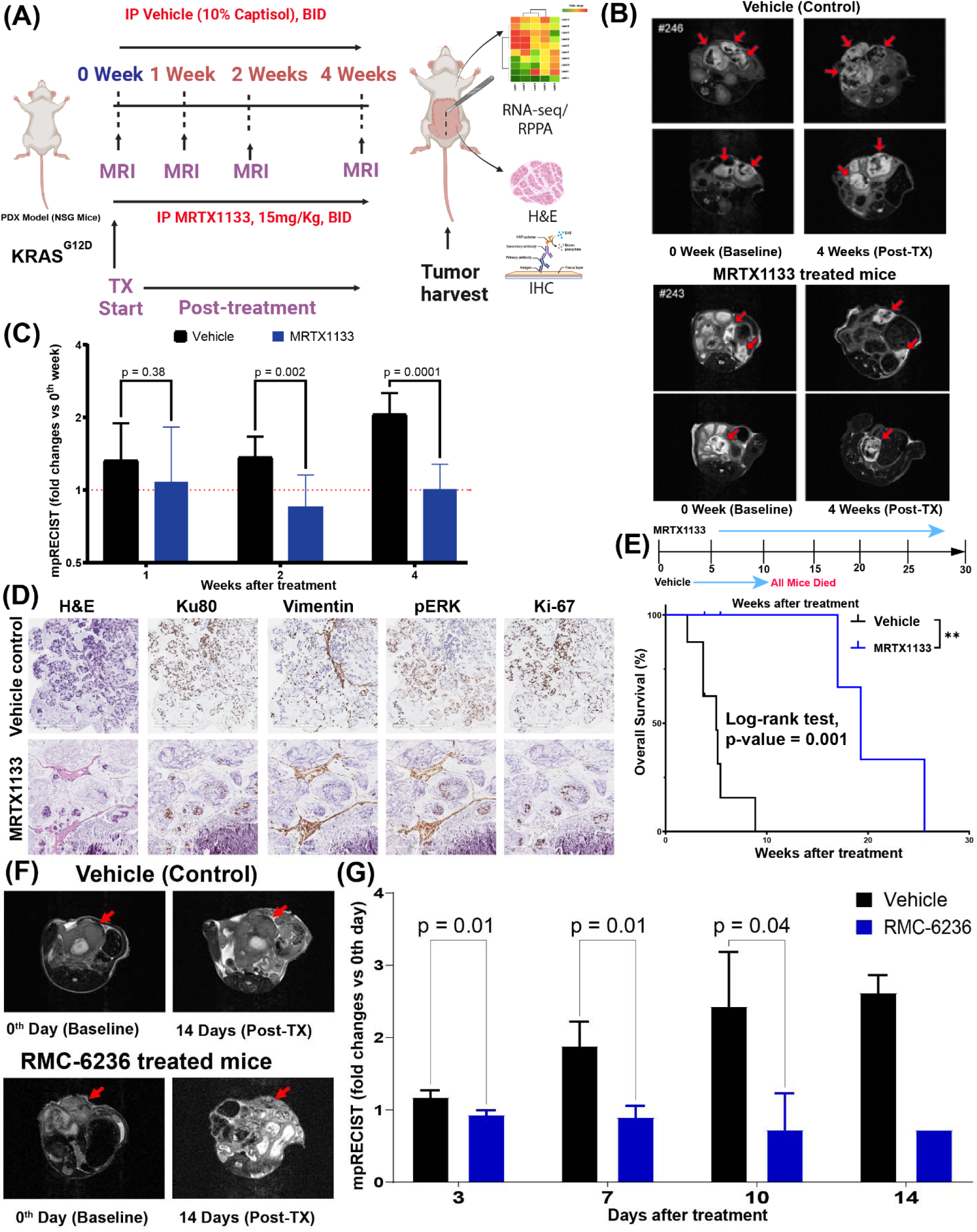
KRAS inhibition is active in PDX models of AA. (A) Schematic representation of PDX experiments conducted using an animal model generated from an AA tumor harboring the *KRAS^G12D^*mutation. (B) MRIs of mice at baseline and after 4 weeks of treatment in vehicle control and MRTX1133-treated mice. (C) Tumor growths after 1, 2, and 4 weeks of treatment in PDX models were evaluated using the mpRECIST method. N=8 in the vehicle control group and n=7 in the MRTX1133 group. (D) H&E and immunohistochemistry (IHC) images of Ku80, Vimentin, pERK, and Ki-67 in PDX tumors from control and treatment groups. (E) Kaplan-Meier plot illustrating the overall survival differences between vehicle control and MRTX1133-treated animals. (F) MRI scans of PDX tumors at baseline and after 14 days of treatment in vehicle control and RMC-6236 treated mice. (G) Fold changes of tumor volume changes measured with respect to the 0^th^ day of treatment in PDX models by mpRECIST for vehicle control (n = 8) vs treatment (n = 7) groups.

### Transcriptional and proteomic response to KRAS inhibition in Appendix Cancer

After confirming the potent anti-proliferative effects of both KRAS inhibitors in organoid and orthotopic PDX models, we performed differential gene expression analyses on RNA-seq from PDX samples (comparing vehicle control vs treatments) to delineate tumor-intrinsic and stromal transcriptional responses to KRAS blockade (**Supplemental Figure 3A and 3D**). Gene Set Enrichment Analysis (GSEA) of human-aligned reads from PDX tumors, representing the tumor transcriptome, revealed significant suppression of the newly published gastrointestinal cancer-specific KRAS-dependent RAS-ERK-mediated upregulated gene set (*20*) (median-rank siKRAS-ERKi-KRASi UP signature) (**Supplemental Figure 3B, C, E, F**) as well as E2F targets, G2M checkpoint genesets (related to cellular proliferation) after treatment with either the pan-KRAS or G12D-specific inhibitor (**Figure 4A**). Key individual genes in the leading edge of the KRAS/ERK signature included *MYC*, *AREG*, and *CCND1* (**Supplemental Figure 3B**). GSEA of the mouse-aligned reads representing the Tumor Microenvironment (TME) showed consistent upregulation of both the alpha and gamma interferon response with both RMC-6236 and MRTX1133 (**Figure 4B**).

**Figure 4:**
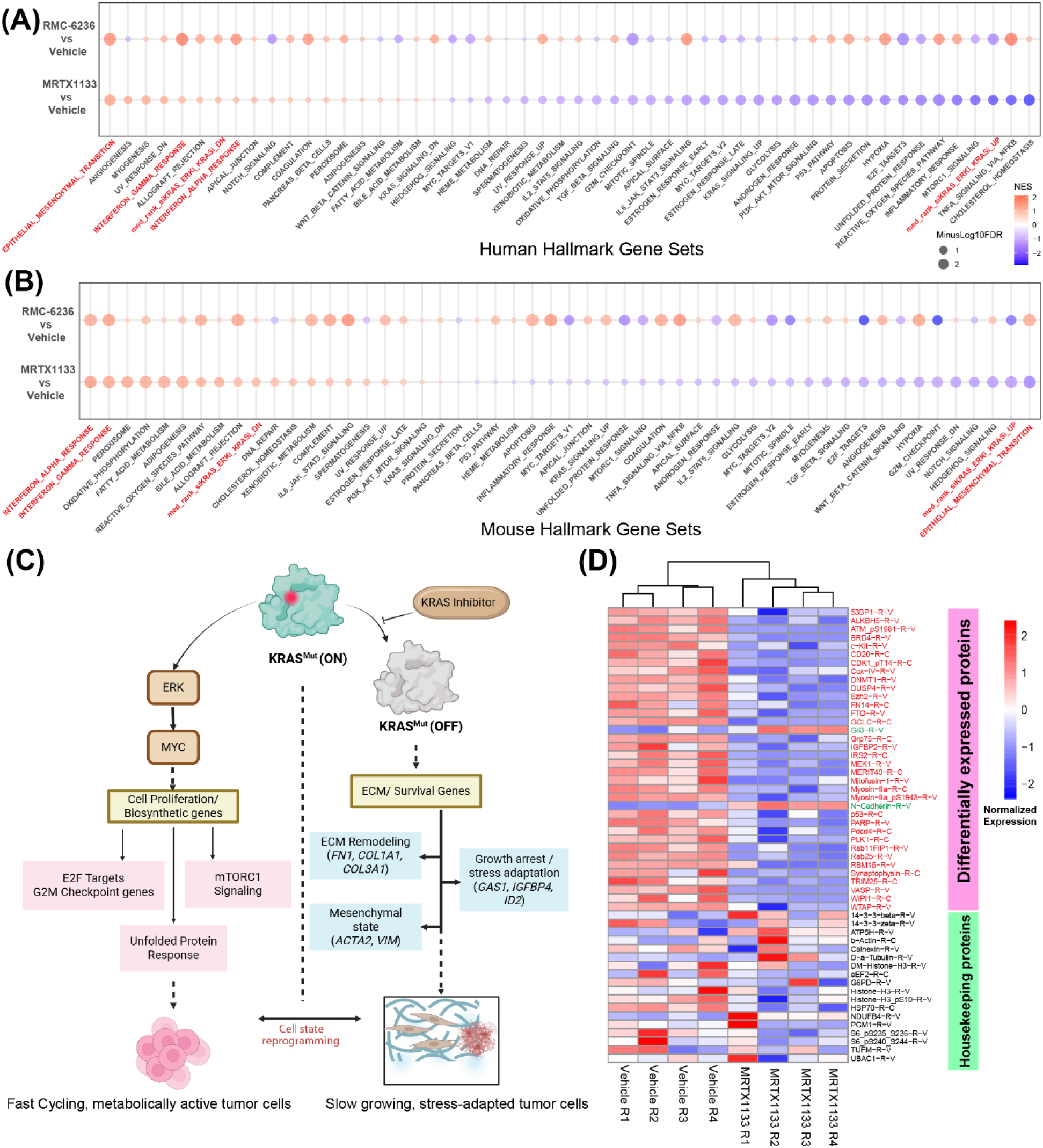
Bulk transcriptional and proteomic analyses of KRAS inhibition. (A) Gene Set Enrichment Analysis (GSEA) of human transcripts (tumor) comparing samples treated with KRAS inhibitor vs. vehicle control. Note downregulation of siKRAS_ERKi_KRASi_Up pathway and upregulation of EMT and interferon gamma and alpha pathways. (B) GSEA of mouse transcripts (stroma) comparing samples treated with KRAS inhibitor vs. vehicle control. Note upregulation of EMT and interferon, interferon gamma, and alpha pathways, and downregulation of siKRAS_ERKi_KRASi_Up pathway. EMT is upregulated in the stroma following pan-KRAS inhibitor RMC-6236 but downregulated following G12D inhibitor MRTX1133. (C) Pathway mechanisms showing cell state transition from proliferating tumor cell state to slow-growing, stress-adapted state after KRAS inhibition. (D) RPPA results comparing MRTX133-treated tumors vs. vehicle control. Note decreases in MAPK activity marker DUSP4 and proliferation marker PLK1, and upregulation of the mesenchymal marker N-Cadherin protein.

The most upregulated pathways in tumor cells included Epithelial-to-Mesenchymal Transition (EMT) and alpha- and gamma-interferon responses, suggesting that these are tumor-intrinsic mechanisms of resistance to KRAS inhibition (**Figure 4A**). While each inhibitor induced a broader EMT-associated transcriptional response, intersection analysis identified a conserved set of 18 leading-edge genes that were consistently upregulated across both KRAS mutant models following treatment. These shared genes (*GAS1, EMP3, LOXL1, BASP1, COL6A3, LRP1, MMP3, COL1A1, COL3A1, COL4A1, COL12A1, VIM, FN1, ACTA2, THBS1, FSTL1, IGFBP4, and ID2*) are functionally enriched for extracellular matrix (ECM) organization, collagen deposition, matrix remodeling, cytoskeletal reprogramming, and growth arrest (**Supplemental Figure 3G, 3H, and 3I**). Notably, the recurrent induction of multiple fibrillar collagens (*COL1A1, COL3A1, COL4A1, COL6A3, COL12A1*), the matricellular proteins *FN1* and *THBS1*, and the collagen-modifying enzyme *LOXL1* indicates a coordinated ECM remodeling response rather than isolated EMT gene activation. Concomitant upregulation of *ACTA2* and *VIM* suggests engagement of a mesenchymal or fibroblast-like transcriptional state, while induction of *MMP3* supports active matrix turnover. In parallel, increased expression of growth-regulatory genes (*GAS1, IGFBP4*, and *ID2*) indicates a shift away from KRAS-driven proliferative programs toward a stress-adapted cellular state following KRAS pathway suppression. Collectively, these data demonstrate that pharmacologic KRAS inhibition—irrespective of KRAS allele specificity—elicits a conserved EMT-linked stromal and extracellular matrix remodeling program in vivo (**Figure 4C**).

Consistent with its known suppression of alpha and gamma interferon, Myc signaling was also downregulated following KRAS inhibition in tumor cells. Interestingly, TNFα signaling via NFkB and cholesterol homeostasis were strongly downregulated by MRTX1133 but not by RMC-6236. Additionally, there was downregulation of Myc, Wnt/B-catenin, E2F, and G2M gene sets, suggesting reduced proliferation in the TME and tumor cells. These transcriptional findings were confirmed by Reverse-Phase Proteomic Array (RPPA) analysis, which showed decreased levels of multiple Ras pathway proteins, including DUSP4(8), and increased N-Cadherin (a marker of EMT), as well as decreased PLK1, a known cell cycle regulator (9) (**Figure 4D**).

### Single-cell transcriptional profiling of tumor cells following KRAS inhibition

In order to better evaluate the differential effects of KRAS inhibition on tumor and TME cells, scRNAseq was performed on AAPDX-01 PDX tumors after 1 week of treatment with MRTX1133. Nearly all tumor cells were killed by MRTX1133 treatment, accounting for only 1.5% of the total cells, compared with 38.3% in tumors treated with vehicle (**Figure 5A**). UMAP clustering of tumor cells revealed clusters aligning with previously identified intestinal epithelial subtypes, including standard differentiated enterocytes, Goblet cells, as well as less differentiated cell types classified as either Transit Amplifying (TA), SERPINE1^high^-EMT, or Nr4a1^high^ cells (**Figure 5B, Supplemental Figure 4A**)(*21–24*). Treated and untreated tumor cells were separated into distinct clusters (**Figure 5C**). DGE analyses followed by GSEA revealed marked downregulation of cholesterol homeostasis, hypoxia, MTORC, and glycolysis pathways, as well as the KRAS-ERK signaling pathway, in treated tumors (**Figure 5D**). Consistent with bulk transcriptomics and RPPA analyses (**Figure 4**), EMT, angiogenesis, myogenesis, and the alpha and gamma interferon pathways were also upregulated in the remaining MRTX1133-treated tumor cells, suggesting that these pathways could all contribute to resistance. Pseudo-time analysis was performed to evaluate the relationship between epithelial cell types, revealing a transition from TA or Nr4a1high cells (enterocytes, goblet cells, or SERPINE1^high^-EMT cells); however, the transition to SERPINE1^high^-EMT cells was not observed in the MRTX1133-treated tumor (**Figure 5E**). These data suggest that KRAS inhibition may limit tumor cells’ ability to undergo EMT.

**Figure 5:**
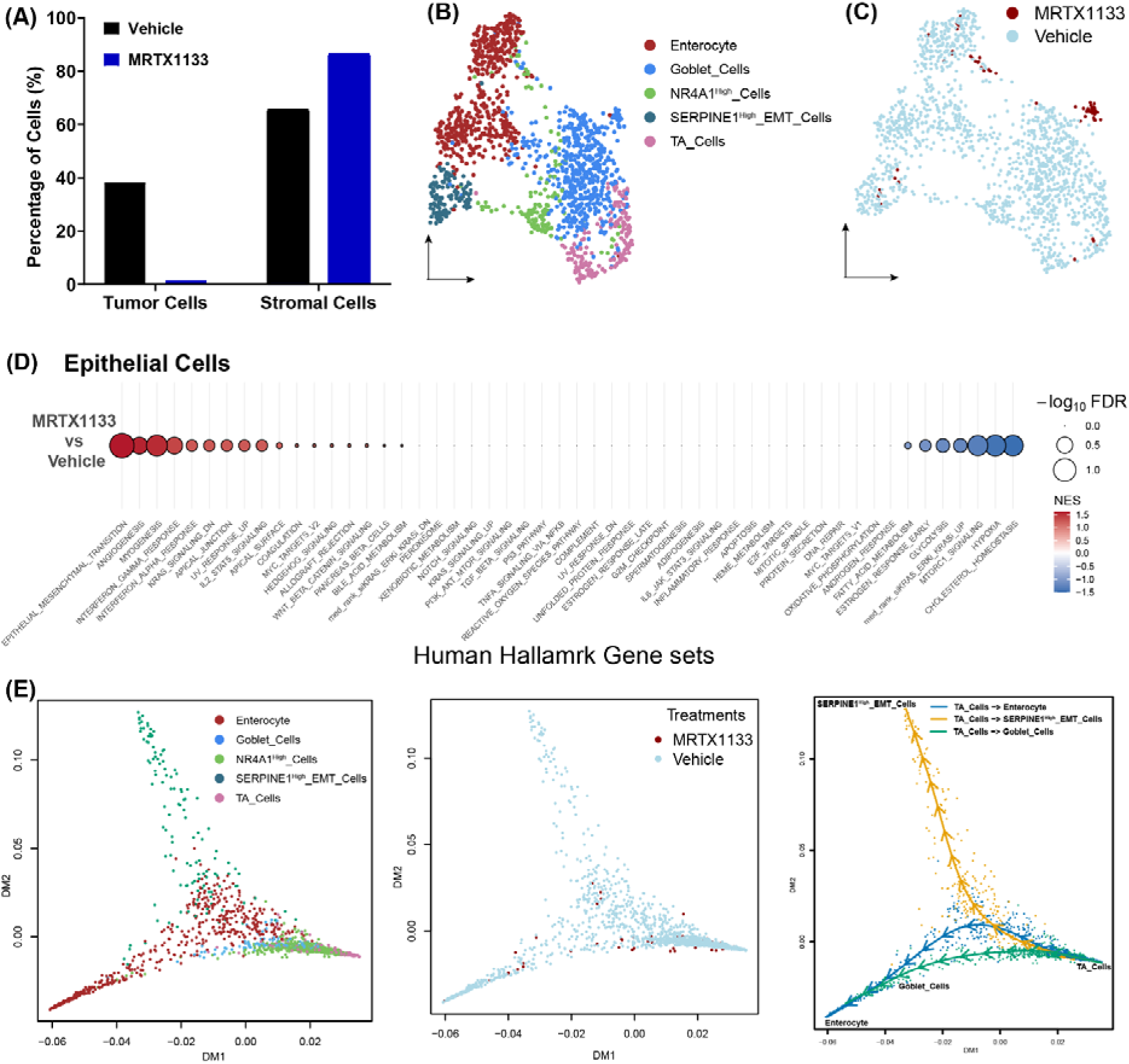
Single Cell RNAseq analysis of tumor cells after KRAS inhibition. (A) Proportions of tumor and stromal cells following MRTX1133 treatment. (B) UMAP plot of tumor cells colored by epithelial cell types. (C) UMAP plot of tumor cells colored by treatment with MRTX1133 or vehicle control. (D) GSEA analysis of tumor cells only, note decreases in cholesterol homeostasis, hypoxia, MTORC, and glycolysis pathways in addition to KRAS signaling and increases in EMT, angiogenesis, myogenesis, as well as alpha and gamma interferon pathways. (E) Pseudo-time analysis identifies a transition from Transit Amplifying (TA) and NR4A1^high^ to either enterocyte, goblet, or SERPINE1^high^ -EMT cells; the transition to SERPINE1^high^-EMT cells is absent in MRTX1133-treated cells.

### Single-cell transcriptional profiling of Tumor Microenvironment cells following KRAS inhibition

TME cells clustered into known cell types, with the first major division between stromal, further subdivided into pericytes, fibroblasts, endothelial, and mesothelial cells, and immune, further subdivided into neutrophils and macrophages (**Figure 6A**, **B, C**). KRAS inhibition had marked effects on the TME, with treated cells clustering separately from untreated (**Figure 6A**). In terms of cell numbers, mesothelial cells were markedly depleted in MRTX1133-treated cells (0.9% vs 17.3%), and fibroblasts were increased (42.2% vs 8.3%; **Figure 6D**). There were also notable changes in the fibroblast population with adventitial fibroblasts (Dpep1^+^, Pdfgra^+^)(*25*), the normal resident fibroblasts in the outer layer of blood vessels (adventitial) and peritoneal spaces, essentially absent, and increased numbers of iCAF in MRTX1133-treated tumors **Figure 6E, F**)(*26–28*). MRTX1133 treatment also led to a greater proportion of INHBA^high^ CAF, which have been implicated in promoting tumor growth, metastasis, and immunosuppression by generating an inflammatory TME (*29, 30*). Pseudo-time analysis of the fibroblast compartment showed a branching trajectory structure with adventitial fibroblasts positioned at the root point and three inferred paths extending toward myCAF, INHBA^high^ CAF, and iCAF endpoints. When stratified by treatment, cells from vehicle control treatment largely occupy the adventitial→myCAF/INHBA-high arms, whereas the iCAF trajectory is exclusively populated by MRTX1133-treated cells, indicating a treatment-associated expansion of the iCAF state rather than a uniform shift across all CAF programs (**Figure 6G**).

**Figure 6:**
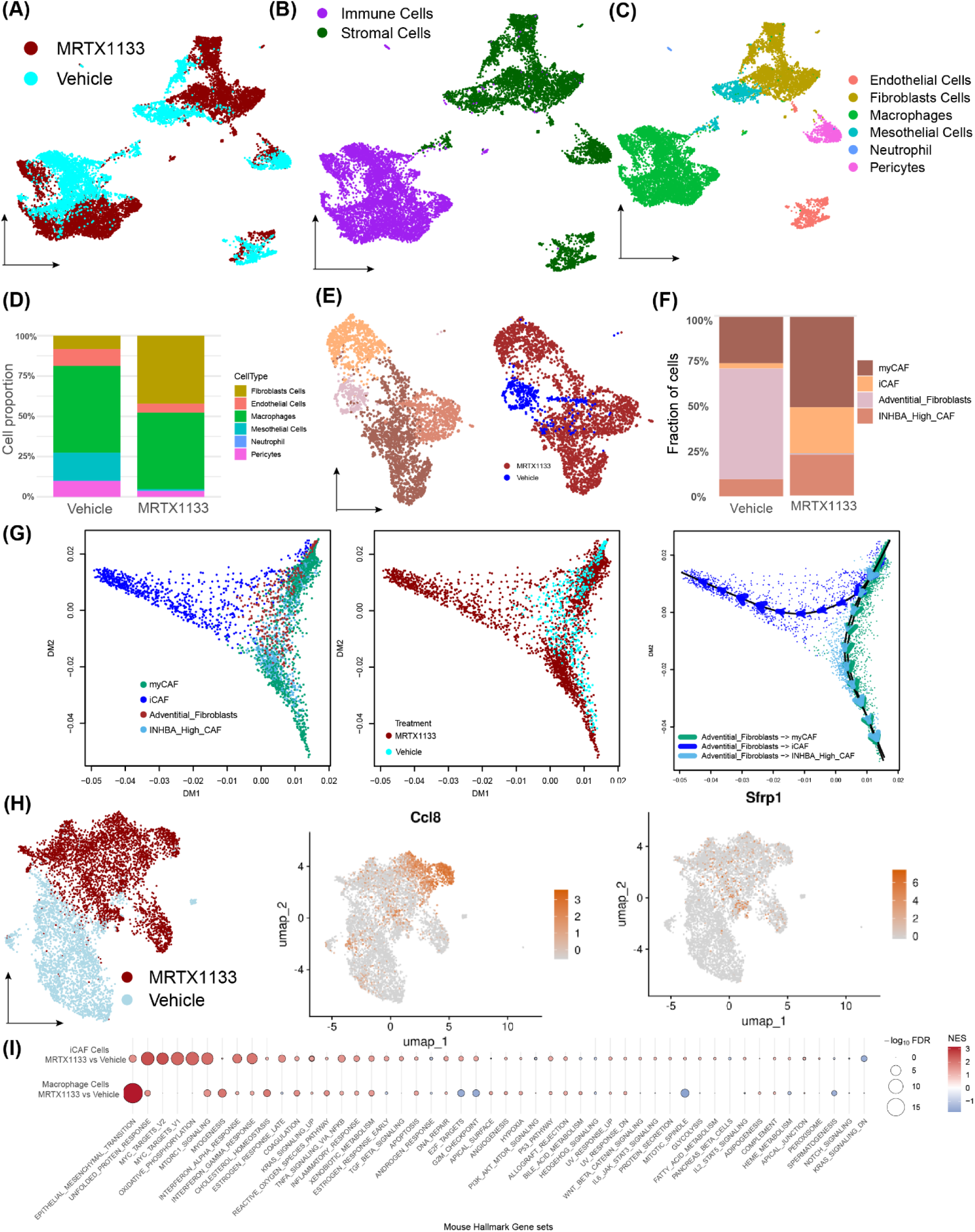
Single Cell RNAseq analysis of stromal and immune cells after KRAS inhibition. UMAP plot of stromal & immune cells colored by (A) treatment, (B) stromal vs. immune lineage, and (C) major cell types. (D) Proportions of different stromal cells after MRTX1133 or control treatment. (E) UMAP plot of fibroblasts colored by fibroblast cell subtypes (left) or treatment (right). (F) Proportions of fibroblasts after MRTX1133 or control treatment. (G) Pseudo-time analysis identifies a transition from adventitial fibroblasts to either myCAF, iCAF, or INHBA^high^ CAF, which occurs primarily in MRTX1133-treated tumors. (H) UMAP plot of macrophages colored by treatment (left), CCL8 expression (middle), or SFRP1 expression (right). (I) GSEA analysis comparing MRTX1133-treated iCAF and macrophage cells to controls; note upregulation of Myc and unfolded protein response in iCAF, and EMT in macrophages.

The macrophage compartment also showed clear separation between treated and untreated samples (**Figure 6H**). In the treated samples, a subpopulation of macrophages upregulated CCL8. CCL8 secreted by macrophages has previously been reported to promote both stemness and invasion via ERK signaling (*31*). A distinct subpopulation of macrophages upregulated Secreted frizzled Related Protein 1 (*SFRP1*), another secreted protein which is known to be a negative regulator of the Wnt pathway(*32*). *SFRP1*-expressing macrophages have been linked to fibrosis in multiple organs (21, 22) and, in one report, to cancer recurrence (*32*). In the GSEA of MRTX1133 vs vehicle control, the iCAF compartment showed broad positive enrichments across multiple hallmark programs, with the strongest signals observed in Epithelial–Mesenchymal Transition (EMT), Unfolded Protein Response, IFN-γ response, and MYC targets (v2), with additional positive enrichment extending into pathways such as oxidative phosphorylation and mTORC1 signaling (**Figure 6I**). In contrast, macrophages showed a different pattern: while EMT is also strongly positively enriched, several cell-cycle/proliferation–linked programs are negatively enriched (blue), including E2F targets, the G2M checkpoint, and mitotic spindle, which trended negative. Overall, the GSEA indicates that KRAS^G12D^ inhibition by MRTX1133 is associated with a strong induction of inflammatory/stress and EMT-like transcriptional programs in iCAFs, whereas macrophages show EMT-like enrichment but a concurrent suppression of proliferative/cell-cycle hallmarks under treatment. These stromal changes in response to a KRAS^G12D^ inhibitor suggest that inhibition of the KRAS/MAPK pathway in tumor cells profoundly influences tumor-cell interactions with the surrounding stroma.

### KRAS inhibition is effective for patients with Appendiceal Adenocarcinoma

Six patients with appendiceal adenocarcinoma treated with a variety of KRAS drugs, including one G12D, two pan-KRAS, and three G12C-specific inhibitors, were identified retrospectively (**Supplemental Table 1, Figure 7**). The cohort included patients with low- and high-grade tumors; most were heavily pre-treated with an average of 3 prior lines of treatment (range 0-6), although one patient was previously untreated. All 6 patients had a biochemical response, as assessed by CEA/Ca19-9 or ctDNA (**Figure 7).** By imaging using standard RECIST 1.1 criteria, which do not necessarily work well for mucinous appendiceal tumors (*2*), all patients showed clinical benefit with 1 CR, 2 PR, and 4 SD for an objective response rate of 50%. Notably, patient #1, with a grade 3 mucinous adenocarcinoma (Muc Adeno) with *KRAS^G12D^* and *GNAS^R201C^* treated on trial with a selective KRAS^G12D^ inhibitor, had a complete response that is ongoing (**Figure 7A, F**). Patient #2 had a grade 2 Muc Adeno with *KRAS^G12C^* treated with Adagrasib, a selective KRAS^G12C^ inhibitor, with >50% decrease in CEA and SD by RECIST criteria, and remains on treatment (**Figure 7B, F**). Patient #3 had a grade 2 colonic-type AA with *KRAS^G12V^* and *TP53^R273H^* mutations, treated on trial with a pan-KRAS inhibitor as 7th-line therapy, achieving SD and a 88.5% decrease in ctDNA while on treatment (**Figure 7C, F**). Patient #4 had a grade 3 colonic-type AA with *KRAS^G12C^*, *TP53*, and *SMAD4* mutations treated on trial with sotorasib, a selective KRAS^G12C^ inhibitor, and Panitumumab, an anti-EGFR antibody. Patient #4’s CEA decreased by 92.0% and achieved PR by RECIST (**Figure 7D, F**). A fifth patient with grade 1 Muc Adeno harboring *KRAS^G12V^*, *GNAS^R201C^*, and PIK3CA*^E545K^*, treated on trial with a pan-Ras inhibitor as 7th-line therapy, has had a 66% drop in Ca19-9 and SD by RECIST after just 2 months of treatment (**Figure 7E, F**). Patient #6 was treated on study with sotorasib and later continued off study after a treatment break with the best response of SD by RECIST (**Figure 7F**).

**Figure 7:**
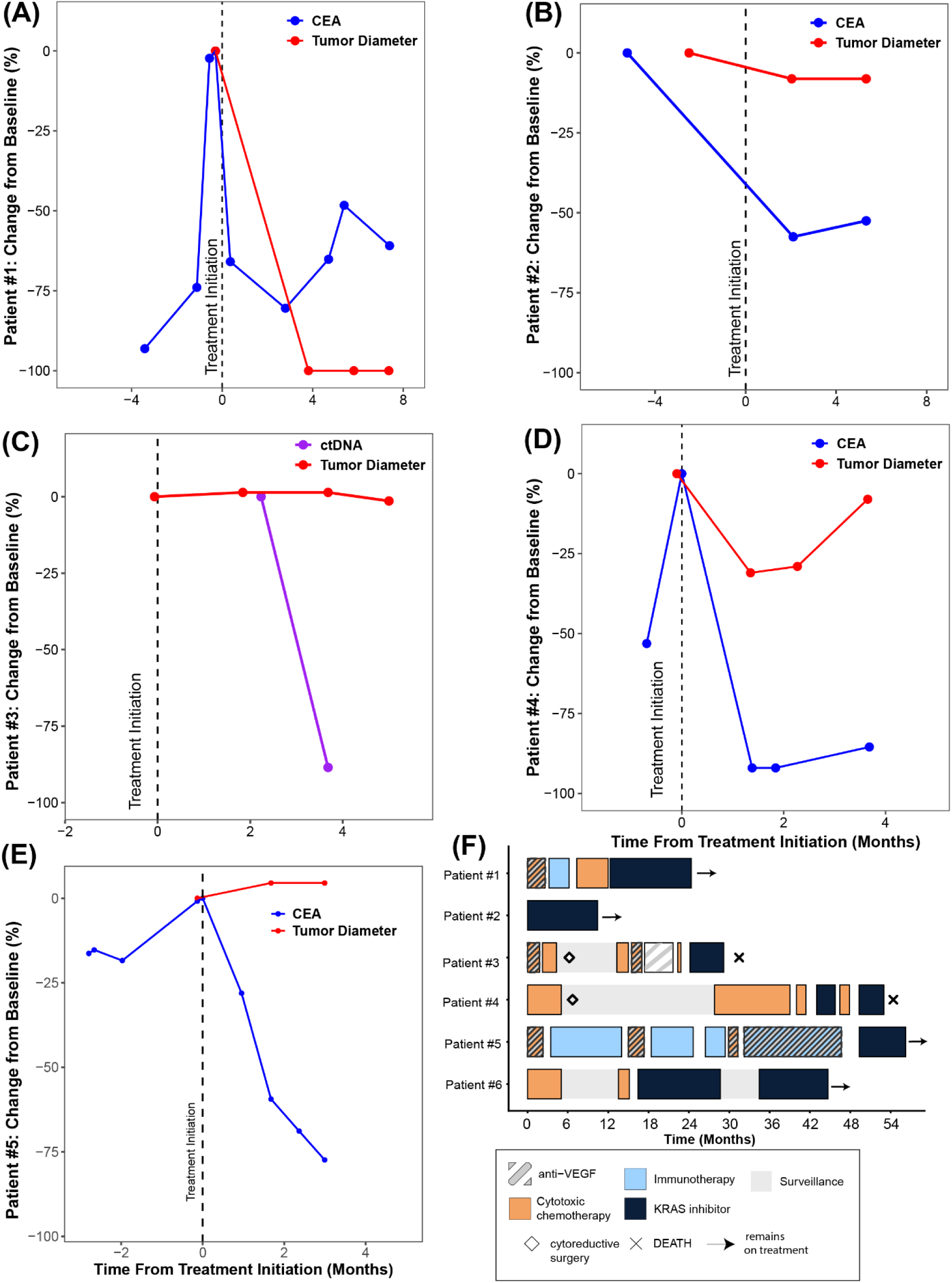
Clinical response to KRAS inhibition in patients with Appendiceal Adenocarcinoma (AA). (A-E) Serum tumor marker or ctDNA and imaging response for patients 1-5. (F) Swimmer plot showing treatment history of six patients with AA treated with KRAS inhibitors, note therapy is ongoing for four patients, indicated by arrow.

## Discussion

Appendiceal cancer, unfortunately, remains an orphan disease with no treatments approved by the FDA specifically for use in appendix cancer. In particular, it has been shown that although commonly used to treat patients with AA, 5FU-based chemotherapy is inactive in low-grade mucinous AA(*2*). Here, we demonstrate that KRAS inhibition is active in preclinical models of AA and in six patients with AA. Given the current lack of effective treatment options for these patients, KRAS inhibition is a promising therapeutic strategy, and there is an urgent need for clinical trials of KRAS inhibitors specifically for patients with KRAS-mutant appendiceal adenocarcinoma. Regarding the mechanism of *KRAS* oncogenesis in appendix cancer, we confirm that, as in PDAC and CRC, AA is KRAS-addicted, with inhibition resulting in decreased cell proliferation and increased apoptotic cell death. Also, similar to PDAC and CRC, the KRAS/MAPK signaling appears to depend primarily on ERK, rather than AKT, as evidenced by the strong alignment with the CRC/PDAC-derived siKRAS-ERKi-KRASi gene signature (*20*). Regarding the comparison between G12D-specific and pan-KRAS inhibitors, we found that in the KRASG12D model, the G12D-specific inhibitor was more potent than the pan-KRAS inhibitor. Although KRAS inhibition was highly effective in these *KRAS*-mutant AA models, we did observe the emergence of resistance, with upregulation of EMT and alpha and gamma interferon responses as potential resistance mechanisms. Interaction between the tumor and the TME also appears to play an important role in the dynamics of KRAS inhibition in AA, with KRAS inhibition altering the TME in a manner that may promote the activity of immune oncology agents. We also identified stromal-mediated mechanisms of resistance, especially in the macrophage compartment, highlighting the complexity of the tumor-TME interaction.

Given the recent report of combined VEGF and PD-L1 inhibition in AA (23), these data suggest promise for combining KRAS inhibition with immune therapy in AA, as is now being pursued in PDAC and other tumor types (*33, 34*). Currently, there are too few AA patients and preclinical models treated with KRAS inhibitors to identify biomarkers of resistance or sensitivity, but notably, responses were observed in both high- and low-grade tumors and in tumors with concurrent *GNAS^R201C^*and *TP53* mutations. Both the PDX and human data also highlight the limitations of standard RECIST criteria when applied to mucinous, peritoneal tumors. Microscopic and molecular studies confirmed that while KRAS inhibition dramatically reduced the number of tumor cells, the change in tumor size measured by cross-sectional imaging (CT or MRI) was modest, given that much of the tumor volume is acellular mucin. Alternate methods to objectively measure tumor response, potentially using serum tumor markers as has been done in ovarian cancer(*35*), or using pathologic criteria(*36*), need to be explored to help facilitate clinical investigation for appendix cancer. Limitations of this study include the limited number of preclinical AA models available for testing, as well as the small sample size and heterogeneous nature of the clinical cohort.

## Materials and Methods

### Generation and expansion of patient-derived organoids

Organoids were generated from each tumor. Fresh tissue samples were placed into separate 50 mL Falcon tubes containing PBS and then transferred to a 10 cm dish. A tissue size of 1 gram was ideal to move forward with processing. Tissues were minced and transferred to a C-tube (Miltenyi Biotec, San Diego, CA, USA) into which 10 mL of tumor digestion media, DMEM medium containing 250 µl FBS (HyClone, Wilmington, DE, USA), 50 mg collagenase II (Sigma-Aldrich, St. Louis, MO, USA), 10 mg dispase II (Sigma), 100 µl penicillin and streptomycin (Sigma), was added. The tissues were dissociated in a fully automated way by using the gentleMACS™ Octo Dissociator with Heaters (Miltenyi Biotec). Fragments were allowed to settle under normal gravity for 1 min and the supernatant was passed through 70 µm cell strainer. Red blood cells in the suspension were lysed with RBC lysis buffer (Roche, Basel, Switzerland) followed by passing through 40 µm cell strainer. The single cell pellets after the centrifugation were then re-suspended in ice-cold growth factor reduced Matrigel. One drop of 50 μl mixture per well were loaded onto 24-well culture plate and allowed to solidify on an incubator for 15 min at 37°C, 5% CO^2^, then 0.5 ml of pre-warmed IntestiCult^TM^ Organoid Growth Medium (STEMCELL Technologies, Vancouver, BC, Canada) was added per well and incubated at 37°C, 5% CO^2^. The growth and quantity of the organoid cultures were monitored. Medium was changed every 2 or 3 days. For drug testing, single cells were obtained from growing Organoids in Matrigel domes using TrypLE Express (Gibco, Grand Island, NY, USA) and suspended in ice-cold Matrigel. Five µl containing 10^4^ cells per well were loaded onto 96-well culture plate and cells were pre-cultured in IntestiCult^TM^ Organoid Growth Medium for 3 days. To test the efficacy of MRTX1133 and RMC-6236, the growth of organoids in the presence of various doses was monitored with Incucyte SX5 (Sartorius, Bohemia, NY, USA) and quantified Organoid Object Total Area (µm²/Image).

### Generation of PDX models

Female NOD.Cg-Prkdcscid Il2rgtm1Wjl/SzJ (NSG) mice, 5-7 weeks of age, were purchased from The Jackson Laboratory (Bar Harbor, ME, USA), and used for tumor implantation. The animals were maintained under specific pathogen-free conditions. Housing and all procedures involving animals were performed according to protocols approved by the Institutional Animal Care and Use Committee of MD Anderson Cancer Center and conducted in accordance with U.S. Common Rule. AA tumors from The Jackson Laboratory for TM00351 and the University of Texas MD Anderson Cancer Center for AAPDX-16 were implanted into NSG mice in the peritoneal cavity. The well-being of the mice was carefully monitored, and mice were sacrificed when signs of disease, mainly abdominal distension together with a rough coat and reduced mobility, were seen. The tumor was harvested, cut into ∼ 25 - 30 mm^3^ fragments, six of which were placed in the peritoneal cavity, both upper and lower abdominal quadrants, and both flank sides, of NSG mice.

### Histological and Immunohistochemistry Analysis

Histological and immunohistochemical analyses were performed in the Department of Veterinary Medicine and Surgery core at MD Anderson Cancer Center. For immunohistochemistry (IHC), we simultaneously assessed multiple protein markers within the paraffin sections. The staining procedure involved the application of multiple primary antibodies, anti-CDX2 (Abcam, EPR2764, Cat# ab76541), anti-KU80 (Cell Signaling, C48E7, Cat# 2180), anti-Vimentin (Cell Signaling, D21H3, Cat# 5741), anti-Ki-67(Abcam, SP6, Cat# 16667), anti-cleaved caspase 3 (Biocare Medical, CP229 a, b, c), anti-Ki-67 (abcam, SP6, Cat# 16667), and anti-phospho-p44/42 MAPK (Erk1/2) (Thr202/Tyr204) (Cell Signaling, D13.14.4E, Cat# 4370), were used. Subsequently, a series of secondary antibodies were utilized to enable the visualization and quantification of distinct protein markers. The quantified data were obtained from at least 4 ROIs per slide in 3 different tissues in each group.

### Drug treatment

MRTX1133 (synthesized in in TRACTION, the University of Texas MD Anderson Cancer Center) were freshly dissolved in 10% Captisol in 50 mM citrate buffer pH 5.0. PDX mice were intraperitoneally (IP) injected with MRTX1133 (15 mg/kg), twice a day. RMC-6236 (Selleck Chemicals, Houston, TX, USA) were dissolved in DMSO, PEG300, and Tween80, then added with ddH_2_O to adjust volume to 0.5 mL per mouse (final concentration of 5%DMSO/40%PEG300/5%Tween/50%ddH2O). PDX mice were IP injected with RMC-6236 (20 mg/kg, an equal mole of MRTX1133) daily. As a control, mice were IP injected with the same amount of vehicle at the same schedule. The IP tumor was visualized using a 7T small-animal MRI system (Bruker Biospin MRI, Billerica, MA, USA), with 35-40 0.75-mm-thick sections, 0.25-mm gaps between sections, throughout the entire peritoneal cavity in each mouse. Tumor growth was evaluated using the modified Peritoneal Response Evaluation Criteria in Solid Tumors (mpRECIST), a quantitative measurement system designed for mucinous peritoneal disease, defined as the sum of the longest diameters of up to 5 target lesions in the abdominal cavity. Tumors were collected for histology, immunohistochemistry, RNA-Seq, and reverse phase protein arrays (RPPA).

### Bulk RNA sequencing

Sample preparation and data preprocessing for bulk RNA sequencing analyses were discussed elsewhere(*37*). Briefly, mRNA was extracted from vehicle- and inhibitor-treated PDX tumors. Human and mouse reads were then separated using Xenome and aligned to their respective reference genomes. The resulting gene-expression counts were used for differential expression analysis and GSEA. Human and mouse hallmark gene sets were used in their respective GSEA analyses.

### Reverse Phase Protein Array

PDX tissue samples were sent to the Functional Proteomics Reverse Phase Protein Array (RPPA) Core facility of the University of Texas MD Anderson Cancer Center. Level 4 (L4) data, which was adjusted for Set-to-Set differences (batch effects) by normalizing identical control samples in the current dataset with an invariant control sample set and applying the adjustment (differences in means × inverse of standard deviation ratio) to each corresponding data point.

### Single-cell RNA sequencing

PDX samples were sent to the Single Cell Genomics Core (SCGC) of the University of Texas MD Anderson Cancer Center. Briefly, 0x Genomics single-cell 5’ R2-only v3 chemistry was used to prepare libraries for paired-end single-cell RNA sequencing. Cellranger (version 8.0.1) was used to convert raw sequence data to FASTQ files. The GRCh38 and mm10 reference genomes were used to align single-cell reads, producing a pair of scRNA-sequencing files (human and mouse) for each tissue sample. scRNA-seq files aligned to human and mouse reference genomes were analyzed separately for downstream analyses.

Seurat (version 5.0.2) was used for quality control, cell filtering, sample integration, unsupervised clustering, and cell-type annotation. For each sample, filtered feature–barcode matrices were imported using the Read10X function, and Seurat objects were constructed with a minimum threshold of 3 cells per gene and 100 detected genes per cell. Cells expressing mitochondrial and ribosomal genes were first verified using gene-name pattern matching. The proportion of mitochondrial and ribosomal transcripts per cell was calculated using PercentageFeatureSet and added to the Seurat object metadata. Quality control metrics were visualized by scatter plots comparing total UMI counts (nCount_RNA) with mitochondrial percentage, ribosomal percentage, and the number of detected genes (nFeature_RNA). Cells were retained for downstream analyses if they expressed more than 200 and fewer than 10,000 genes, had less than 20% mitochondrial transcripts, and less than 40% ribosomal transcripts.

Filtered cells were normalized using log normalization (LogNormalize) with a scaling factor of 10,000. Highly variable genes (n = 2,000) were identified using the variance-stabilizing transformation (VST) method. Gene expression values were scaled across all genes while regressing out total UMI counts, mitochondrial gene percentage, and ribosomal gene percentage. Principal component analysis (PCA) was performed using the highly variable genes, and the top 100 principal components were computed. For downstream analyses, the first 50 principal components were used to construct a shared nearest neighbor (SNN) graph. Cell clustering was performed using the Louvain algorithm across multiple resolutions ranging from 0.1 to 1.0. Uniform Manifold Approximation and Projection (UMAP) was used to visualize the data in two dimensions, with a random-seed control to ensure reproducibility. Putative doublets were identified using the DoubletFinder package. Parameter optimization was conducted using a parameter sweep across the first 50 principal components, followed by selection of the optimal pK value. The expected number of doublets was estimated assuming an 8% doublet rate. Cells classified as singlets were retained for subsequent analyses. To correct for batch effects across samples, Harmony integration was applied using sample identity as the grouping variable. Neighborhood graph construction, clustering across multiple resolutions, and UMAP visualization were repeated using the Harmony-corrected embeddings. For the human reference genome–aligned data, the Harmony-integrated object was used to extract tumor cell populations, based on the biological assumption that human-derived epithelial cells represent malignant tumor cells within PDX samples. Tumor cells were therefore identified and subsetted based on expression of canonical epithelial marker genes (*EPCAM, KRT8*, and *KRT18*). Conversely, for the mouse reference genome–aligned data, the Harmony-integrated object was used to isolate stromal cell populations. Mouse-derived cells lacking expression of epithelial marker genes were considered host-derived stromal cells and were selected for downstream stromal-specific analyses. This dual-genome, species-resolved strategy enabled robust separation of tumor and microenvironmental compartments in PDX models while minimizing cross-species contamination and preserving biologically relevant cellular heterogeneity. Markers used for annotating major subtypes of epithelial, stromal, and fibroblast cells are shown in **Supplementary Figure 4A-C**.

### Clinical Data

Under the guidance of an IRB-approved study with an informed consent waiver, the Foundry software system was used to query the internal MD Anderson appendix cancer database and identify all patients with appendiceal adenocarcinoma treated with a KRAS inhibitor. Charts were manually reviewed by physicians to extract treatment histories. The clinical histories for these patients are summarized in **Supplementary Table 1**.

### Statistical analyses and software

Statistical analysis was performed using GraphPad Prism 10 (GraphPad Software) or R v4.3.2 in RStudio v2023.09.01). One-way ANOVA with the Holm-Šidák post hoc test for multiple comparisons, or the Student t-test for two-group comparisons, was used to assess the significance of differences. When appropriate, we estimated within-group variation and ensured that it was similar across groups being statistically compared. The log-rank test was used to determine statistically significant differences between two Kaplan-Meier survival curves.

## Declarations

### Ethics approval and consent to participate

Patient data were retrospectively collected under the guidance of an IRB-approved study with an informed consent waiver approved by the University of Texas MD Anderson Cancer Center. All animal studies were performed according to protocols approved by the Institutional Animal Care and Use Committee of MD Anderson Cancer Center and conducted in accordance with U.S. Common Rule.

### Consent for publication

Not Applicable

### Availability of data and materials

The datasets generated and/or analysed during the current study will be uploaded to the GEO database, and the accession id will be published in the manuscript.

### Competing interests

Dr. John Paul Shen: Grant/Research support/ Collaboration:

Natera, Inc; BostonGene; Revolution Medicine; Summit Therapeutics

Consulting / Stock ownership: Engine Biosciences, NaDeNo Nanoscience

Patent: Siegel D, Schweer J, Abagyan R, Shen JP. SMALL MOLECULE FOR TREATMENT OF CANCER OF THE APPENDIX. Filed July 17, 2023, PCT/US2023/027939

Dr. David Hong: Grant/Research support/ Collaboration: 280 Bio, AbbVie, Adaptimmune, Adlai-Nortye, Amgen, Astelles, Astra-Zeneca, Bayer, BeiGene USA, BioBridge, Biomea Fusion, Bristol-Myers Squibb, Deciphera, E.R. Squibb & Sons LLC, Eisai, Eli Lilly, Endeavor, Erasca, Exelixis Inc., F. Hoffmann-LaRoche, Genentech, Immunogenesis, Incyte Inc., Merck, Mirati, NCI-CTEP, Novartis, Pfizer, Quanta Therapeutics, Revolution Medicines, STCube Pharmaceuticals, VM Oncology, and Yiling Pharmaceutical.

Travel, accommodations, or expense support: American Association of Cancer Research (AACR), American Society of Clinical Oncology (ASCO), Bayer, BeiGene USA Inc., Genmab, Medscape, Mirati Therapeutics Inc., Pfizer, Society for the Immunotherapy of Cancer (SITC), and Telperian.

Consulting, speaker, or advisory role: 280 Bio, Acuta Capital Partners LLC, Alpha Insights, Amgen, Bayer, Boxer Capital, Children’s Oncology Group, COR2ed, Cowen Group Inc, Crossbridge Bio, Ecor1 Capital, Erasca, Gerson Lehrman Group Inc., Group H, Guidepoint, Immunogenesis, Jansen Pharmaceuticals, Kestrel Therapeutics, Medacorp, Medscape, Orbi Capital, Pfizer, Revolution Medicines, T-Knife, Travistock Group, WebMD, and Yiling Pharmaceutical.

Ownership interests: Molecular Match (Advisor), OncoResponse (Founder, Advisor), Telperian (Founder, Advisor), and CrossBridge Bio (Advisor).

Dr. Michael J. Overman: Consulting/advisory role: Bristol Myers Squibb, Roche/Genentech, Gritstone Bio, MedImmune, Novartis, Array BioPharma, Janssen, Pfizer, 3D Medicines, Merck, Eisai, Simcere;

Research funding: Bristol Myers Squibb, Merck, Roche, MedImmune.

### Funding

This work was supported by the Col. Daniel Connelly Memorial Fund, the Andrew Sabin Family Fellowship Award (J.P.S. is an Andrew Sabin Family Foundation Fellow at The University of Texas MD Anderson Cancer Center), the National Cancer Institute Cancer Center Support Grant (P30 CA016672), the Cancer Prevention & Research Institute of Texas (RR180035 & RP240392 to J.P.S., J.P.S. is a CPRIT Scholar in Cancer Research), the Appendiceal Cancer Pseudomyxoma Peritonei Research Foundation (Catalyst Research Grant to J.P.S.), and a Conquer Cancer Career Development Award (2022CDA-7604125121 to J.P.S.). Any opinions, findings, and conclusions expressed in this material are those of the author(s) and do not necessarily reflect those of the American Society of Clinical Oncology or Conquer Cancer.

### Authors’ contributions

S.C and I.I. contributed equally. S.C, I.I., and J.P.S. wrote the main manuscript text. J.P.S. conceptualized and designed the whole experiment. S.C. and I.I. performed experiments, data acquisition, and analyses. V.P. and S.K. prepared Figure 7. V.P., S.K., A.Y., M.Y., N.H., M.O., and D.H. helped obtain clinical samples for organoid and PDX models and extract treatment response data from the institutional database. J.P.S., V.P., S.K., B.H., K.F.F., A.N.S., and M.J.O. analyzed and interpreted the patient data. M.W.T. and W.C.F. performed all histopathological experiments and related data analyses of this work. All authors reviewed the manuscript.

## Acknowledgements

The authors acknowledge the support of the High Performance Computing for research facility at the University of Texas MD Anderson Cancer Center for providing computational resources that have contributed to the research results reported in this paper. The authors thank the CPRIT Single Cell Genomics Core (RP180684) for support with single-cell sequencing experiments and the Reverse Phase Protein Array (RPPA) Core facility at MD Anderson.

## Supplemental Figures & Legends

**Supplemental Figure 1:**
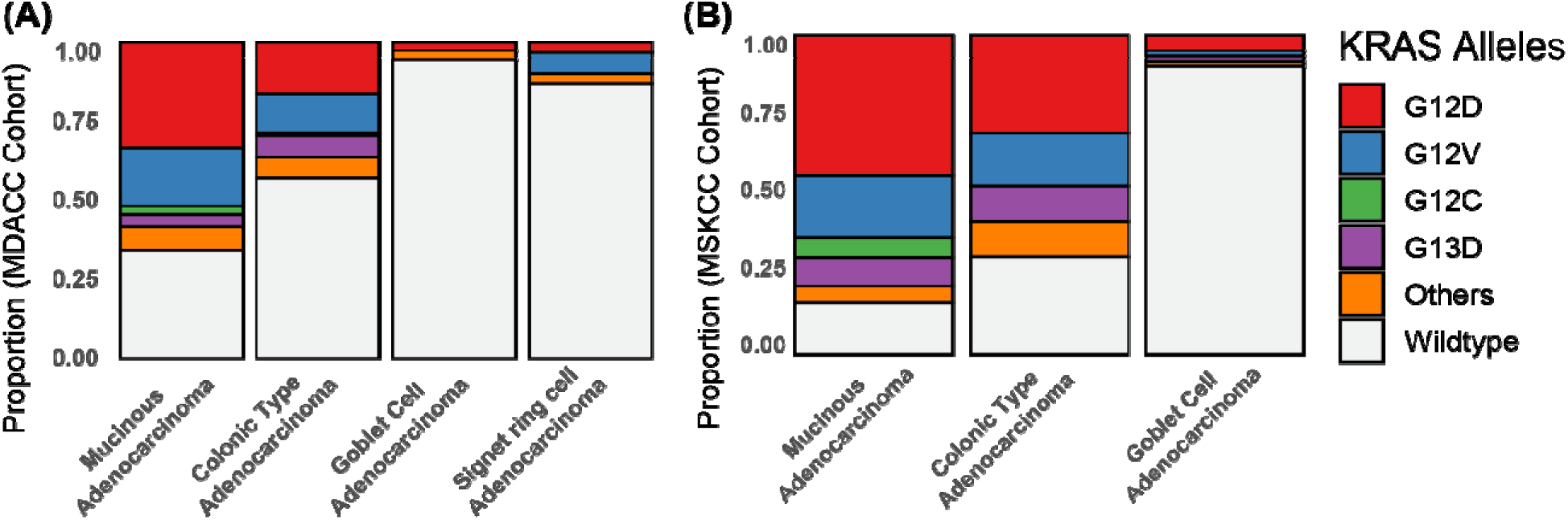
KRAS mutant allele frequency. Distributions of different *KRAS* alleles across different tumor histologies in (A) MDACC and (B) MSKCC cohorts.

**Supplemental Figure 2:**
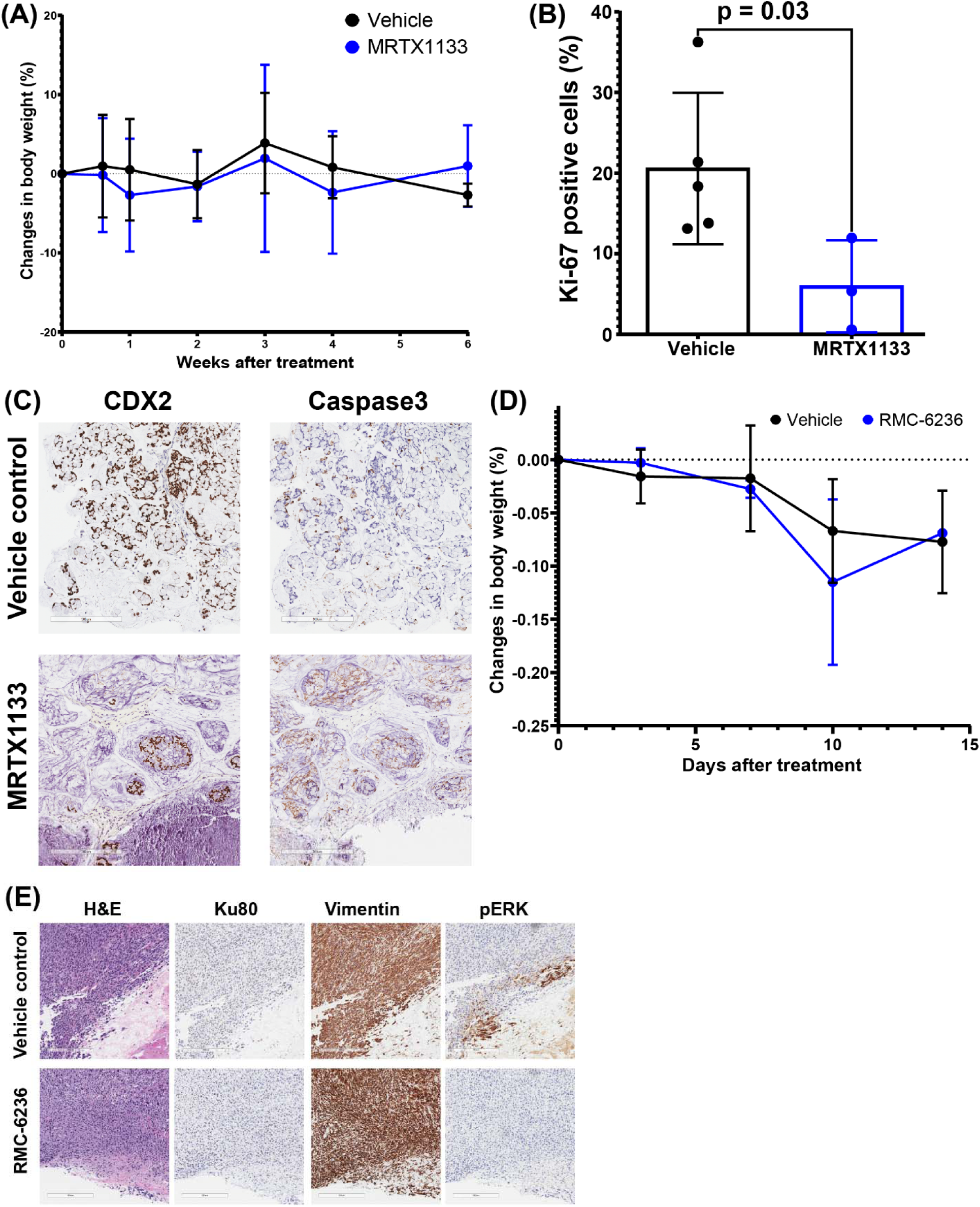
Effects of KRAS inhibition *in vivo*. (A) Change in body weight for mice treated with MRTX1133 or control. (B) Fraction of cells positive for Ki67 following treatment with MRTX1133 or control. (C) CDX2 and Caspase3 staining after treatment with MRTX1133 or vehicle control, note the decrease in CDX2+ (tumor) cells and increase in caspase3 specifically in CDX2+ cells. (D) Change in body weight for mice treated with RMC-6236 or vehicle control. (E) Ku80 (marker of human cells), vimentin (marker of fibroblasts), and pERK IHC following treatment with RMC-6236 or vehicle control, note reduced presence of Ku80+ (human tumor) cells and decreased pERK following RMC-6236 treatment.

**Supplemental Figure 3:**
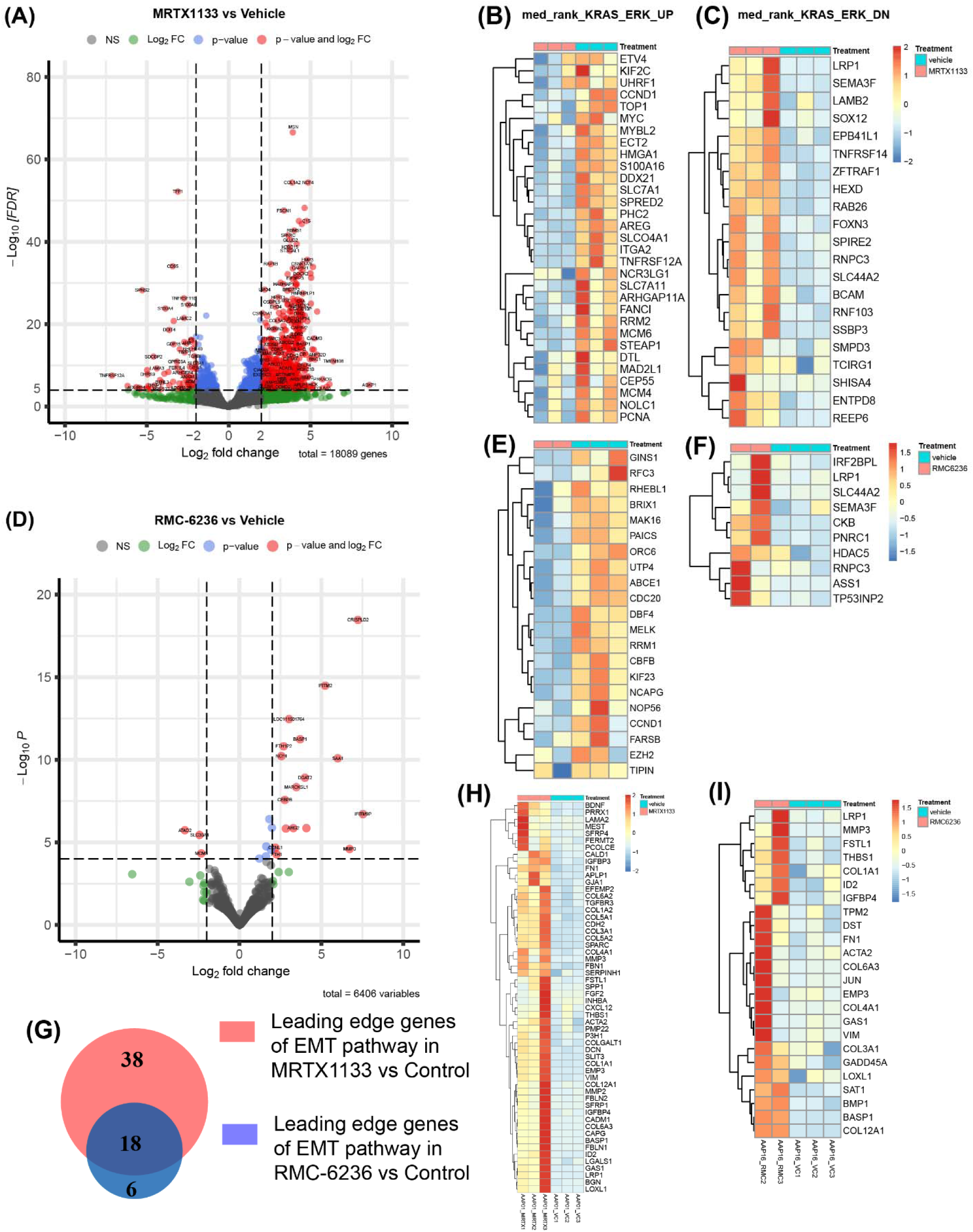
Transcriptional effects of KRAS inhibition *in vivo*. (A) Volcano plots from differential gene expression analyses performed between MRTX1133-treated tumors and vehicle control. Leading edge genes contributing enrichment scores of (B) KRAS-ERK_UP and (C) KRAS-ERK_Down gene sets in GSEA of MRTX1133 vs Control analysis. Note decreased expression of *MYC*, *AREG*, and *CCND1* following MRTX1133 treatment. (B) Volcano plots from differential gene expression analyses comparing RMC-6236-treated tumors to vehicle control. Leading edge genes contributing enrichment scores of (E) KRAS-ERK_UP and (F) KRAS-ERK_Down gene sets in GSEA of RMC-6236 treated tumors vs vehicle control. (G) A Venn diagram showing common and unique genes identified in the leading-edge genes of EMT pathway observed in GSEA analyses of MRTX1133-treatment vs vehicle control and RMC-6236-treatment vs vehicle control. Leading edge genes contributing enrichment scores of the EMT pathway in GSEA analyses of (I) MRTX1133-treatment vs vehicle, and (J) RMC-6236-treatment vs vehicle control.

**Supplemental Figure 4:**
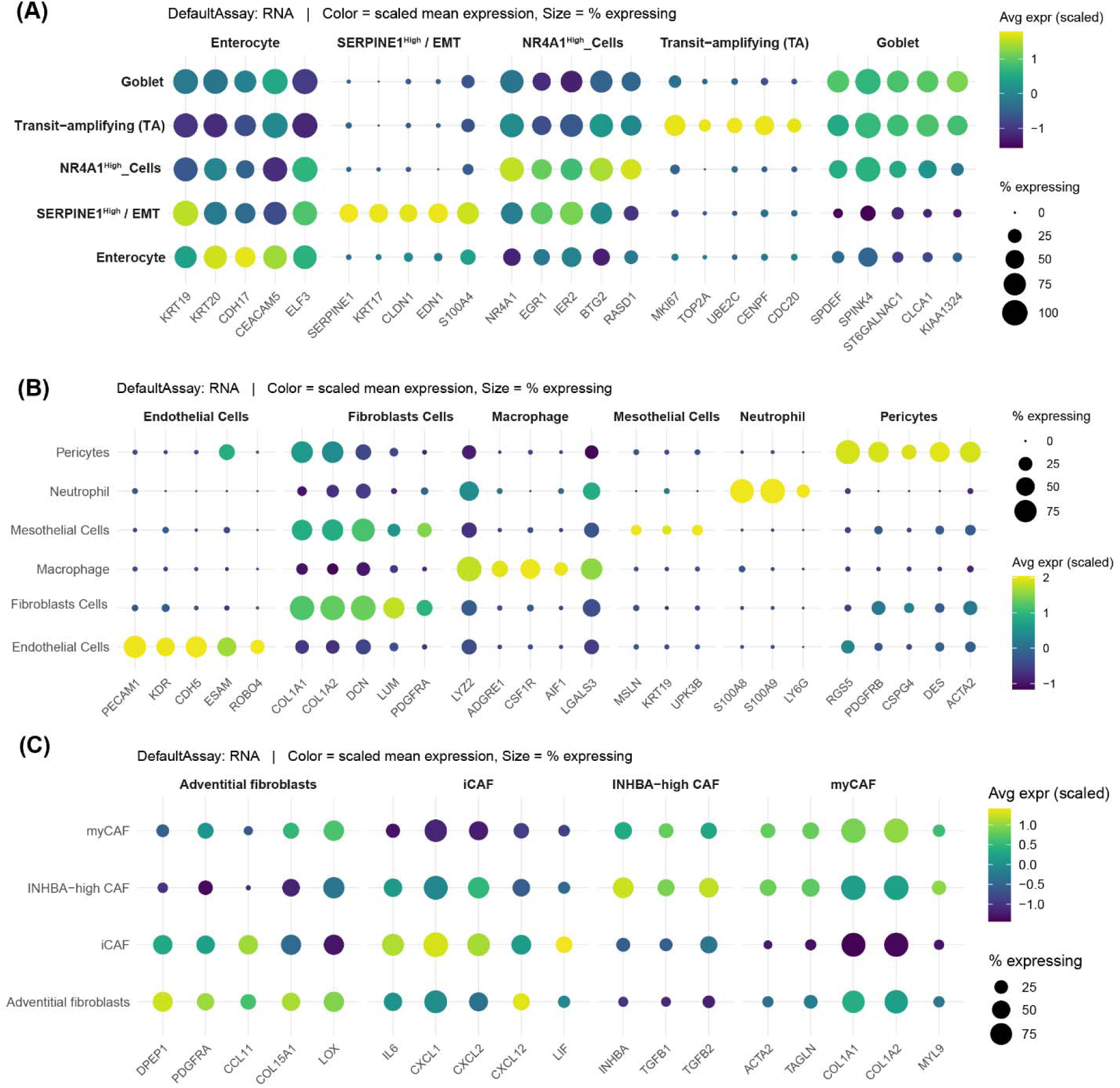
Markers for epithelial, stromal, and fibroblast cell subtypes. Dot plot identifying major subtypes of (A) epithelial cells, (B) Stromal cells, and (C) Fibroblast cells.

**Supplemental Table 1:** Summary information for clinical cohort.

